# Immunological landscape of human lymphoid explants during measles virus infection

**DOI:** 10.1101/2022.09.12.507535

**Authors:** Joshua A Acklin, Aum R Patel, Andrew P Kurland, Shu Horiuchi, Arianna S Moss, Emma J Degrace, Satoshi Ikegame, Jillian Carmichael, Shreyas Kowdle, Patricia Thibault, Naoko Imai, Hideki Ueno, Benjamin Tweel, Jeffrey R Johnson, Brad R Rosenberg, Benhur Lee, Jean K Lim

**Affiliations:** Department of Microbiology, Icahn School of Medicine at Mount Sinai, New York, NY, United States; Graduate School of Biomedical Sciences, Icahn School of Medicine at Mount Sinai, New York, NY, United States; Department of Otolaryngology, Icahn School of Medicine at Mount Sinai, New York, NY, United States

## Abstract

In humans, lymph nodes are the primary site of measles virus (MeV) replication. To understand the immunological events that occur at this site, we infected human lymphoid tissue explants using a pathogenic strain of MeV that expresses GFP. We found that MeV infected between 5-15% of cells across donors. Using single cell RNA-Seq (scRNA-Seq) and flow cytometry, we found that while most of the 29 cell populations identified in the lymphoid culture were susceptible to MeV, there was a broad preferential infection of B cells and reduced infection of T cells. Further subsetting of T cells revealed that this reduction may be driven by the decreased infection of naïve T cells. Transcriptional changes in infected B cells were dominated by an interferon stimulated gene (ISG) signature. To determine which of these ISGs were most substantial, we evaluated the proteome of MeV-infected Raji cells by mass spectrometry. We found that IFIT1, IFIT2, IFIT3, ISG15, CXCL10, MX2, and XAF1 proteins were the most highly induced, and positively correlated with their expression in the transcriptome. These data provide insight into the immunological events that occur in lymph nodes during infection and may lead to the development of therapeutic interventions.

## INTRODUCTION

Measles virus (MeV) is the most infectious human virus, with a reported R_0_ value of 12-18 (1–7). MeV outbreaks have largely been controlled with the advent of the two-dose measles, mumps, and rubella (MMR) vaccine, yet MeV still causes ∼200,000 deaths annually, primarily among unvaccinated children in developing countries (8–10). However, recent surges in vaccine-hesitancy have allowed MeV to re-emerge in countries like the United States and the United Kingdom, where the MMR vaccine coverage has historically been high (11–14). With many global vaccination campaigns for MeV stalled due to the COVID-19 pandemic, the risk of MeV outbreaks globally continues to grow (15). Compounding these crises is the lack of any licensed antiviral that targets MeV once individuals are infected (16, 17).

MeV is a morbillivirus of the family *Paramyxoviridae* that is transmitted through the respiratory tract, where alveolar macrophages and dendritic cells are the initial cellular targets of infection (18, 19). These infected immune cells then traffic to the draining lymph nodes where the virus replicates rapidly in lymphocytes that express the entry factor CD150/SLAMF1 (20–23), followed by egress through lung epithelium that is mediated by basolateral expression of the Nectin-4 receptor (23–27). MeV is also known for causing immune amnesia through the depletion of CD150^+^ B and T cells in both primary and secondary lymphoid organs, increasing the morbidity and mortality rates from secondary infections with common childhood pathogens (28–35). Immunological amnesia following MeV infection has been shown to markedly reduce the antibody repertoire towards common childhood pathogens, such as the human parainfluenza viruses, respiratory syncytial virus, coronaviruses, and cytomegalovirus (36). Given that immune responses are primarily generated in the draining lymph node, and that this site is a critical launching pad for MeV infections, understanding virus: host interactions at this site is paramount for identifying factors that shape disease progression.

Modern reanalysis of early work on immunological amnesia caused by measles implicates T cells due to a delayed type-I hypersensitivity response to tuberculin antigen (37, 38). In vitro characterization of MeV infection in primary lymphocytes revealed that B cells are the most extensively infected population of lymphocytes, consistent with their high CD150 expression (39). Characterization of MeV infection in PBMCs from both humans and macaques further demonstrated the propensity of MeV for lymphocytes and that MeV infection is biased towards naive B cells, memory B cells, and memory T lymphocytes, which was subsequently validated in PBMCs derived from MeV-infected children (18, 34, 40, 41).

While these studies provide insights into MeV infection of lymphocytes, they do not examine infection within the complex architecture of secondary lymphoid tissue. The draining lymph nodes are organized with high-density B cell follicles surrounded by T cell zones (42), which may be important to determine the in vivo cellular susceptibility as well as the kinetics of lymph node infection. Studies in macaques have examined the geographic distribution of infected cells within secondary lymphoid tissues and have identified that the majority of infection is established within B-cell follicles (41). Further, these studies recapitulated the heightened susceptibility to infection among memory but not naïve lymphocyte subsets within secondary lymphoid organs (41). Studies using human tonsil explants have also been conducted, where lymphoid tissue structures and native cell ratios are intact, which found that B cells are a preferential target of MeV infection and that memory T cells were more extensively infected compared to naive T cell subsets (43, 44). In this study, we revisited this human lymphoid explant model using a GFP-expressing pathogenic isolate of MeV which has been shown to mimic human clinical outcomes in non-human primates, commonly referred to as a “wild-type” isolate (45). Our findings both confirm and extend our understanding of cellular susceptibility to MeV in humans. While a similar analysis of MeV-infected airway epithelium has been conducted (46), we present transcriptional signatures of MeV-infected lymphocytes with single cell resolution in human lymphoid tissue explants and link these transcriptional signatures with translated products in the proteome.

## RESULTS

### MeV replicates efficiently in human lymphoid tissues ex vivo

To evaluate how MeV infection proceeds within human lymphoid tissue, we infected human tonsil tissues ex vivo, the most accessible lymphoid tissue for laboratory use. Tissue samples from routine, non-inflamed tonsillectomies were infected with a pathogenic isolate of MeV (IC323) that expresses GFP (MeV-GFP) as previously described (44, 45, 47). While this model lacks a functional lymphatic system, and thus does not exactly mimic the way that MeV enters the draining lymph node during human infections, it benefits from retaining the 3-dimensional tissue architecture of human lymph tissue. To assess the extent to which MeV could replicate in human lymphoid tissues across eleven donors, we collected culture supernatants at days 3, 6, and 8 post-infection and measured virus production by fluorescence plaque assay. As shown in **Figure 1A**, viral titers increased for all donors, with ∼2.5 log increase over the 8-day culture.

**Figure 1:**
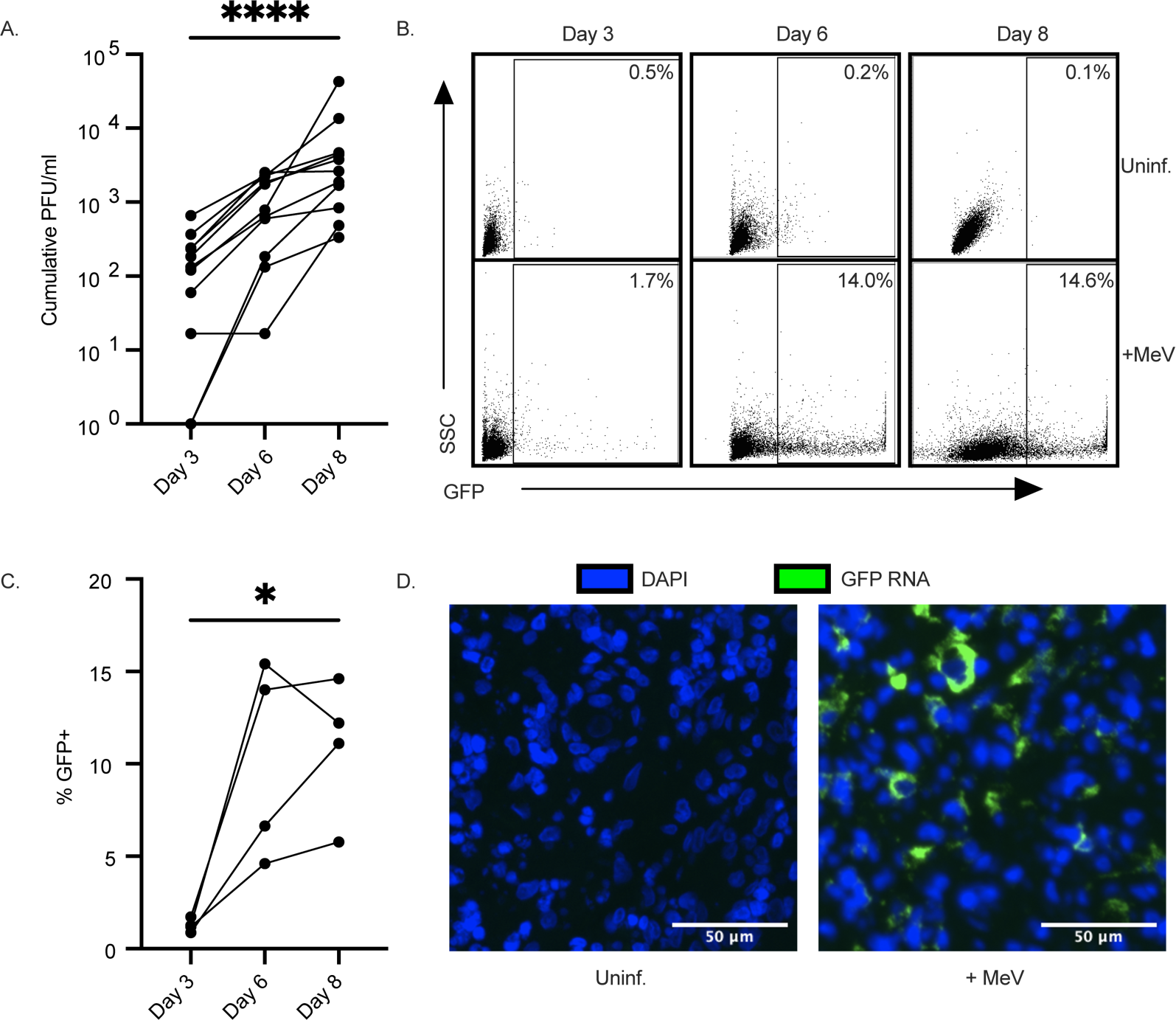
MeV productively infects human lymphoid tissue explants. Tonsil tissues (n=11) were infected with MeV-GFP. Cumulative viral plaque forming units (PFU/ml) was quantified from supernatants collected on days 3, 6, and 8 post-infection by plaque assay **(A)**. Representative flow plots quantifying infection (GFP) are shown for one donor **(B)** and quantified across 4 donors over time **(C)**. In situ hybridization for GFP RNA (green) on paraffin-embedded tissues collected on day 8 post-infection compared to a donor-matched, uninfected control **(D)**. Nuclei were counterstained with DAPI (blue). Scale bars represent 50µm. Significance was determined by one-way ANOVA using Friedman’s test with Dunnett’s multiple comparison test. *indicates *p*<0.05, ** indicates *p*<0.01, *** indicates *p*<0.001 and **** indicates *p*<0.0001.

To assess the extent of infection within the tissues and to further confirm productive infection, we measured the frequency of GFP^+^ cells over time by flow cytometry. As shown in **Figure 1B**, for one representative donor, the percentage of GFP^+^ cells increased over time, while no GFP signal was detected in the uninfected condition. Quantification across 4 donors revealed a frequency of MeV-infected cells that ranged between 5-15% of live cells by day 8 post-infection (**Figure 1C**). Further, RNA transcripts for GFP were readily detected within the tissues through in situ hybridization (**Figure 1D**). Together, our data establish that human tonsils infected ex vivo are susceptible to MeV without any stimulation or infection enhancers, providing us with a robust model system to further characterize MeV-infected cells.

### scRNA-Seq of MeV-infected tissues reveals broad cellular susceptibility

Given the high percentage of GFP^+^ cells detected, we sought to sort and analyze infected cells using scRNA-Seq. To do this, we generated single cell suspensions from MeV-infected and donor-matched uninfected tonsil tissue on day 8 post-infection, the time point at which the maximum number of infected cells was observed. We sorted GFP^+^ (infected) and GFP^-^ (bystander) cells from the MeV-infected condition, as well as the GFP^-^ cells (uninfected) from the uninfected condition for scRNA-Seq using a workflow shown in **Figure 2A**. We validated the quality of sequencing across the three groups by quantifying unique molecular identifier (UMI) counts, unique genes captured, and the representation of mitochondrial transcripts **(Supplementary Figure 1A-C**). As expected, only cells from the infected group had appreciable MeV transcripts (**Figure 2B**). Among the infected cells, we examined the expression levels of viral transcripts as a final confirmation of infection status. Infected cells showed a transcriptional gradient of viral genes from 3’ to 5’, consistent with the phenomenon of run-off transcription that occurs for paramyxoviruses (**Figure 2C**).

**Figure 2:**
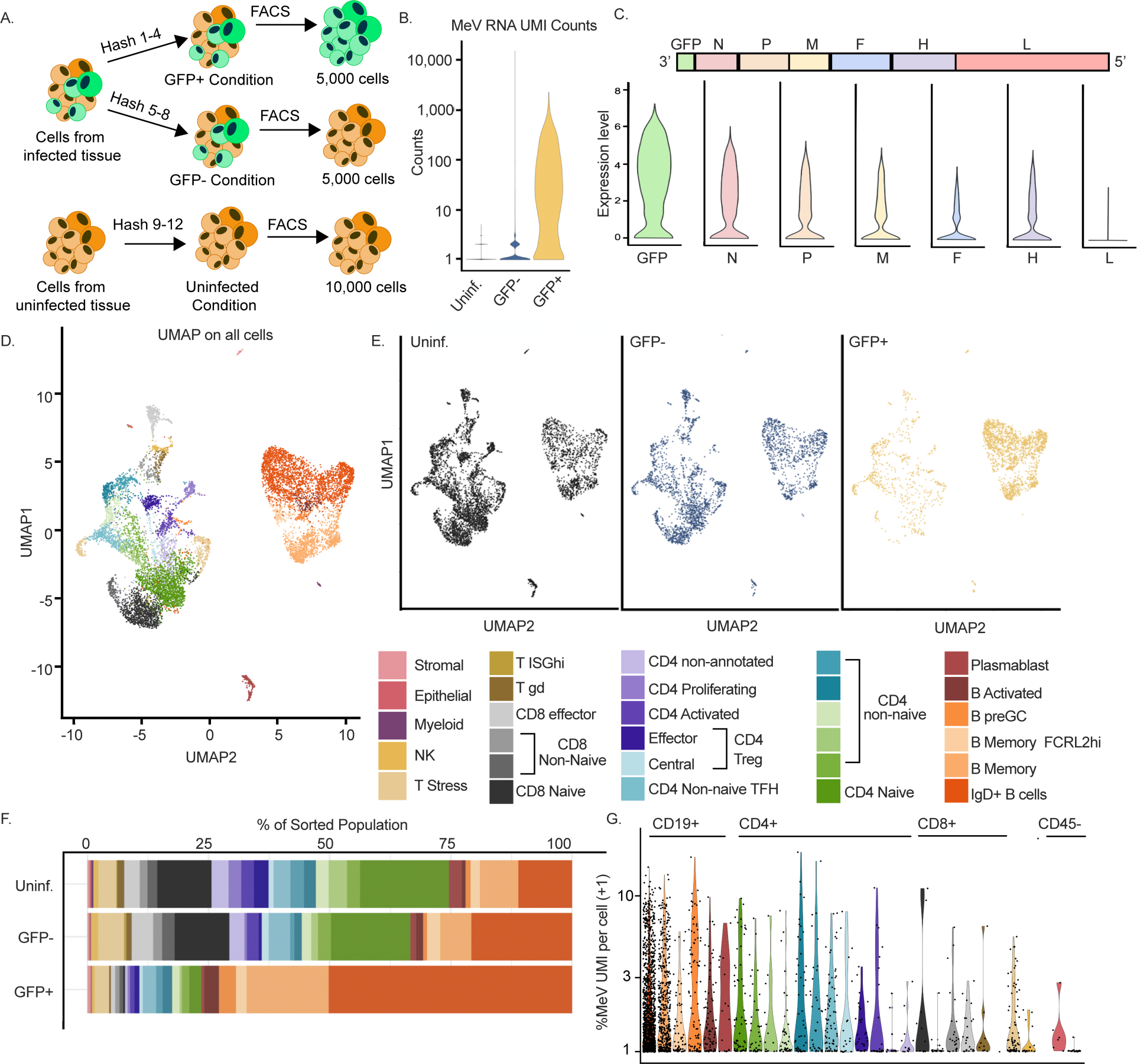
scRNA-Seq identifies 29 unique cell populations in tonsils susceptible to MeV. Tonsil tissue from MeV-GFP-infected and uninfected explants on day 8 from one donor were sorted for scRNA-Seq. Schemata of scRNA-Seq workflow are shown in **(A)**. Cells from the infected condition were sorted and hash-labeled into GFP^+^ and GFP^-^ groups. Uninfected GFP^-^ cells were sorted from a donor-matched uninfected control. 5,000 GFP^+^ cells, 5,000 GFP^-^ cells, and 10,000 uninfected cells were encapsulated for sequencing. MeV RNA unique molecular identifiers (UMIs) were quantified for quality control and filtering **(B)**. Normalized expression of each MeV transcript in infected cells was quantified and shown as violin plots ordered from 3’ to 5’ in the MeV genome **(C)**. Canonical correlation analysis (CCA) was conducted on all groups (combined) and individual clusters were functionally annotated (see also **Figure S1**). Clusters were visualized by UMAP **(D)** and then split into conditions based on captured hashing oligonucleotide sequences for further analysis **(E)**. The frequencies of each cell cluster identified in **(E)** were calculated for each group, and quantification is shown in **(F)**. The percentage of the transcriptome that is MeV RNA is shown in **(G)**, with a +1 pseudocount artificially added to the values for display on a log_10_ axis. All cluster annotations are labeled by the color legend shown.

Following data integration, we annotated constituent cell clusters across conditions based on immune cell reference data and supervised differential marker gene analysis (see *Methods*). Using this strategy, we identified 29 distinct populations of cells (**Figure 2D**), with the vast majority belonging to either T or B cell subsets (annotation strategy shown in **Supplementary Figure 1D**). Interestingly, GFP^+^ transcripts were identified in all 29 populations, albeit to different levels in each population, spanning subpopulations of CD4 and CD8 T cells, B cells, tonsillar epithelium, and tonsillar stromal cells (**Figure 2E**). We were surprised by this finding and assessed the expression of *SLAMF1*, the gene encoding the canonical MeV receptor CD150, in each cluster. As shown in **Figure S1E**, the detection of *SLAMF1* in our dataset was not robust. However, we do detect higher levels among B cell populations, particularly among activated B cells, which is consistent with previous reports (39, 41, 43). We next quantified the frequency of each of these populations within the uninfected, bystander, and infected cell-sorted sample groups and found that, despite T cells comprising the largest population of cells in human tonsils (as shown in uninfected and bystander groups), they were not the majority among the MeV-infected cells. Instead, IgD^+^ B cells were overrepresented within the pool of GFP^+^ cells (**Figure 2F**). To assess the extent to which the transcriptome is co-opted for viral gene expression, we quantified the percentage of UMI counts that mapped to MeV UMIs for each identified cell population. As shown in **Figure 2G**, MeV transcripts constituted ∼0.93% of all transcripts for each cluster on average (calculated using the median % viral UMI per cluster). One possible explanation for differences in the frequency of cell clusters among the GFP^+^ condition would be that cells had either proliferated or died during or because of preferential infection of various cell subsets found within the tissue. To assess this possibility, we evaluated the B cell clusters for gene signatures associated with proliferation. As shown in **Figure S1F-G**, while we observed high pathway scores for S phase and G2/M phase among CD4^+^ Cells (annotated as proliferating in **Figure 2**), we did not observe any elevation within any B cell cluster, regardless of infection status. However, as this analysis was limited to a single donor at a single time point, we can only conclude that MeV has a wide cellular tropism within the lymphoid explants and suggest that IgD^+^ B cells are the primary target.

### B cells are preferential targets of MeV infection in lymphoid tissue explants

While the scRNA-Seq analysis suggested that differences in susceptibility to MeV may exist within the lymphoid tissue explants, these data evaluated only a single time point for a single donor. To define MeV infection across donors and over time, we immunophenotyped major cell subsets identified in the scRNA-Seq dataset by flow cytometry (n=3). We first quantified the frequency of MeV-infected B and T cells and compared these frequencies to their frequency among GFP^-^ bystander cells (from the infected condition) and donor-matched, uninfected cells (gating schemata in **Figure 3A**). For ease of data interpretation, we also show the frequencies of GFP over time within cell populations discussed in this section in **Figure S2** and present a complete gating schema for all analyses in **Figure S3**. As shown in **Figure 3B**, there was a significantly higher frequency of CD19^+^ B cells among the GFP^+^ cells compared to their frequency in the uninfected or bystander populations. This difference was established by day 3 and maintained across the 8-day culture. Conversely, we observed a decrease in the frequency of CD3^+^ T cells (**Figure 3C**). Evaluation of CD150 expression by flow cytometry showed a trend toward higher levels of CD150 on CD19^+^ B cells compared to CD3^+^ T cells. While this did not reach statistical significance, this trend is consistent with the higher CD150 transcript counts observed in B cell populations compared to T cell populations by scRNAseq (**Figure 3D; Supplementary Figure 1E**).

**Figure 3:**
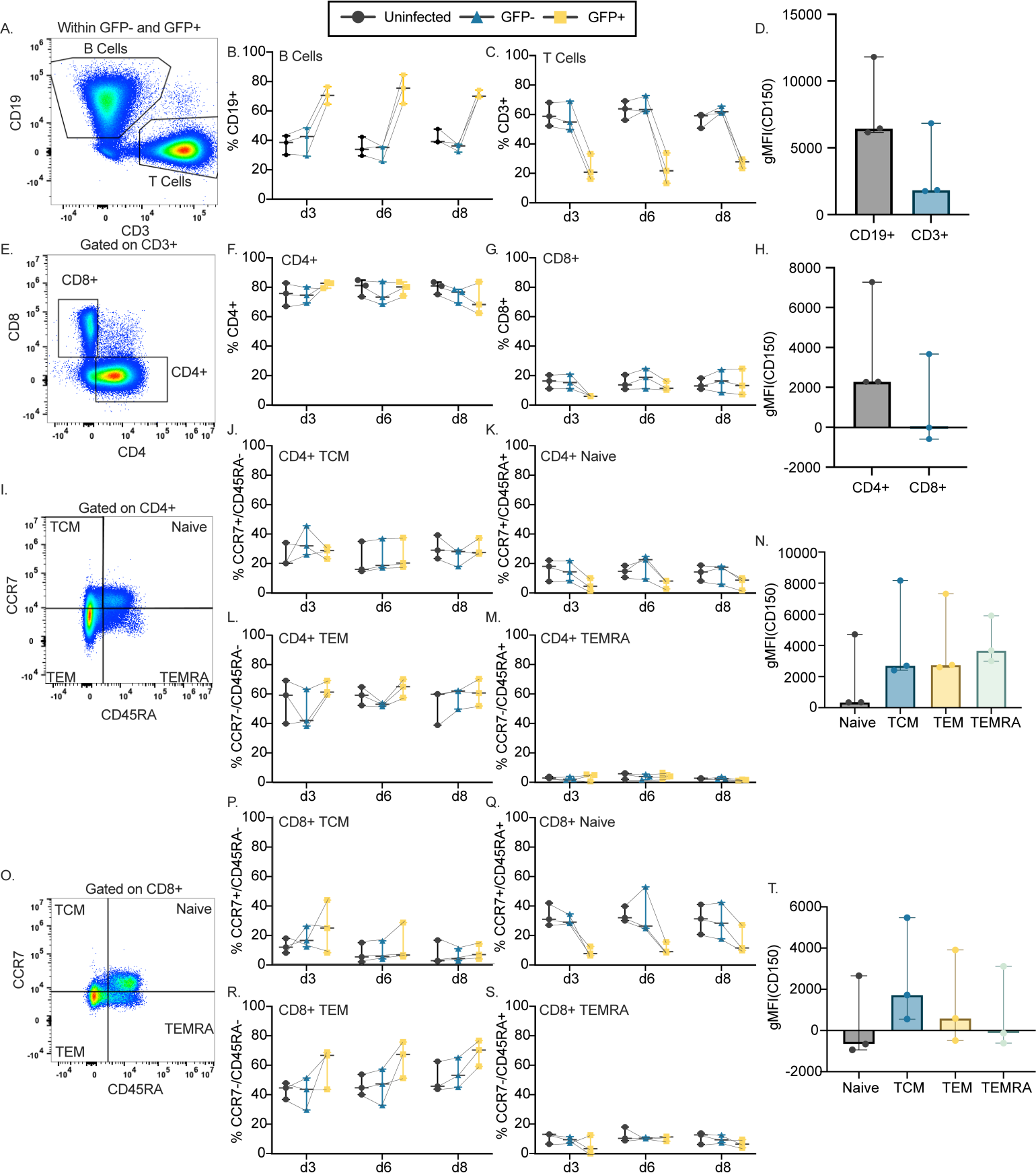
MeV preferentially infects B cells and is restricted among naïve T cells. Cells from MeV-GFP-infected and donor-matched uninfected tissues were collected at days 3, 6, and 8 post-infection, and were then immunophenotyped by flow cytometry (n=3; gating schemata in **A, E, I,** and **O**). The frequency of CD19^+^ B cells **(B)** and CD3^+^ T cells **(C)** among all CD45^+^ cells are quantified and compared over time for GFP^+^ cells, GFP^-^ bystander cells, and donor-matched uninfected controls. The mean fluorescent intensity (MFI) of surface CD150 among CD19^+^ and CD3^+^ cells is compared **(D)**. Susceptibility to infection among CD4 (**F**) and CD8 (**G**) populations is shown, with CD150 expression compared between populations (**H**). Naive (CD45RA^+^/CCR7^+^), TCM (CD45^+^/CCR7^-^), TEM (CD45RA^-^/CCR7^-^), and TEMRA (CD45RA^+^/CCR7^-^) populations are quantified and compared among CD4^+^ cells (**I-M**) and CD8^+^ cells (**O-S**). CD150 expression is compared among CD4^+^ **(N)** and CD8^+^ **(T)** subpopulations. For all immunophenotyping panels, significance was determined by two-way ANOVA using the Geisser-Greenhouse correction with Tukey’s multiple comparison test. For panels (**D**) and (**H**), significance was determined by the Wilcoxon matched-pairs signed rank test. For panels (**N**) and (**T**), Significance was determined by one-way ANOVA using Friedman’s test with Dunnett’s multiple comparison test. For all plots, the median with the 95% confidence interval are shown.

We next asked whether the preference for B cell infection was driven by a specific B cell subset, or if all B cells were more susceptible to infection. To test this, we subset B cells based on CD38 and CD27 expression (gating schemata shown in **Supplementary Figure 4A**). To discriminate between susceptibility differences shared among all B cells and those that are subtype-specific, we calculated the frequency of each B cell subset as a percentage of the total B cell pool among both GFP^+^ B cells from infected tonsil explants, as well as among bystander and uninfected B cells. We found essentially no differences in infection based on CD38/CD27 expression (**Figure S4B-E**). The lack of differences in infection between subsets is also consistent with the relatively stable expression of CD150 among these populations **(Figure S4F)**. Given the dramatic increase in the IgD^+^ B cell cluster observed among GFP^+^ cells in the scRNA-seq **(Figure 2F**), we next assessed if IgD status conferred heightened susceptibility to infection among B cells across time. We recapitulated the finding that IgD^+^ cells are more frequent among the GFP^+^ population than among bystander and uninfected cells (**Figure S4G**). However, when examined among CD19^+^ cells in each group, we found no evidence of preferential infection based on IgD status (**Figure S4H-I**). Likewise, CD150 expression was not different between IgD^+^ and IgD^-^ cells at day 6 (**Figure S4J**). These data show that while B cells are more susceptible to MeV infection, this is most likely not driven by any individual subset of B cells.

We also evaluated the susceptibility of T cell subpopulations by examining the frequency of CD4^+^ and CD8^+^ cells among GFP^+^, bystander, or uninfected cell subsets. As shown in **Figure 3E-G**, we identified no differences in susceptibility between helper (CD4^+^) and cytotoxic (CD8^+^) designations, consistent with the lack of differences in CD150 expression, which was generally low, between CD4^+^ and CD8^+^ T cells (**Figure 3H**).

### Reduced susceptibility of naive T cell subsets in human lymphoid tissue

Previous reports have shown that mature (CD45RA^-^) T cell subsets are more susceptible to MeV infection than naive (CD45RA^+^) T cells (34, 40, 41). To assess this in our model, we evaluated the frequency of naïve T cells (CD45RA^+^CCR7^+^), as well as non-naïve T central memory (TCM; CD45RA^-^CCR7^+^), T effector memory (TEM; CD45RA^-^CCR7^-^) and T effector memory RA^+^ (TEMRA; CD45RA^+^CCR7^-^) subsets among CD4^+^ and CD8^+^ T cells from the GFP^+^, GFP^-^ (bystander) and uninfected groups. As shown through **Figure 3I-T**, we observed reduced MeV infection in CD4^+^ and CD8^+^ naïve T cells compared to the non-naïve subsets, which were less frequent in the GFP^+^ population than in the bystander or uninfected groups, particularly at earlier time points. Evaluation of the non-naïve subsets did not reveal an increased frequency among GFP^+^ cells, suggesting they were no more likely to become infected than their proportion in the culture. Assessment of CD150 expression on these T cell subsets shows that naïve T cells trended toward less CD150 expression than mature subsets, these differences were not significant, suggesting that CD150 expression alone does not explain these trends in susceptibility. Lastly, we hypothesized that the proximity of follicular CD4^+^ T cells to highly susceptible B cells in the follicle could impact the susceptibility of these CD4^+^ cells to infection. To test this, we evaluated the frequency of MeV-infected cells among CD4^+^ CD45RA^-^ T cells based on CXCR5 expression. As shown in **Figure S5**, we found no difference in susceptibility based on CXCR5 status. Taken together, these data indicate a reduced susceptibility of naïve T cells to MeV that is largely independent of CD4 or CD8 status, CD150 expression, and CXCR5 expression.

### MeV induces a canonical ISG response in both B and T cell transcriptomes

Given that the most striking susceptibility differences to MeV infection were observed between B and T cells, we next asked whether the host response to infection among these cell types could contribute to these differences in susceptibility. To test this, we randomly sampled an equal number of B cells or T cells from the uninfected, bystander, and GFP^+^ groups from the scRNA-Seq data and conducted differential gene expression analysis. As expected, the most significantly induced genes among the infected cells were MeV genes and GFP transcripts (**Figure 4A-B**). Following the viral genes, the most significantly upregulated transcripts among GFP^+^ B and T cells were associated with a canonical interferon (IFN) signature. This pattern of interferon induction was strikingly similar among bystander cells, which were GFP^-^ (thus not containing viral transcripts). We detected expression of the edited MeV interferon antagonist transcript, *V*, in GFP^+^ cells but were unable to make meaningful comparisons in the expression of these transcripts among infected cell clusters due to the low read coverage at the p-editing site, where non-templated nucleotide insertion distinguishes *V* transcripts from the more abundant *P* mRNAs (**Supplementary Figure 6**). To directly compare the host response in infected B and T cells, we constructed a Venn-diagram of significantly induced genes in each group. As shown in **Figure 4C**, we find a highly conserved response between both cell types, among which *IFIT1, IFIT2, IFIT3, MX1, MX2*, *XAF1,* and other canonical interferon stimulated genes (ISGs) were shared. While the only gene found to be uniquely induced in infected B cells as compared to infected T cells was the MeV *P* gene, T cells were found to have induced additional ISGs that were not significant in B cells, including *OAS1, OAS2, OAS3*, *OASL*, *USP18*, *HELZ2*, *SAMD9L,* and *HERC6*. This unique pattern of gene expression may be biologically meaningful, or instead a consequence of strict statistical thresholding. Careful and directed comparative analyses using protein-based approaches will need to be conducted to confirm the relevance of these differences. To further validate this IFN signature, we conducted qRT-PCR from the tissues of two additional tonsil donors over time. We evaluated the expression of *IFIT1*, *IFIT3,* and *MX1*, three of the highly expressed type-I interferon stimulated genes (ISGs) from the scRNA-Seq analysis. As shown in **Figure 4D-F**, we found potent induction of all three genes by day 8 post-infection, the time point of scRNA-Seq. Taken together, these data suggest MeV induces a potent interferon response at the transcriptional level in both infected and bystander B and T cells, with no notable differences that account for the increased susceptibility of B cells.

**Figure 4:**
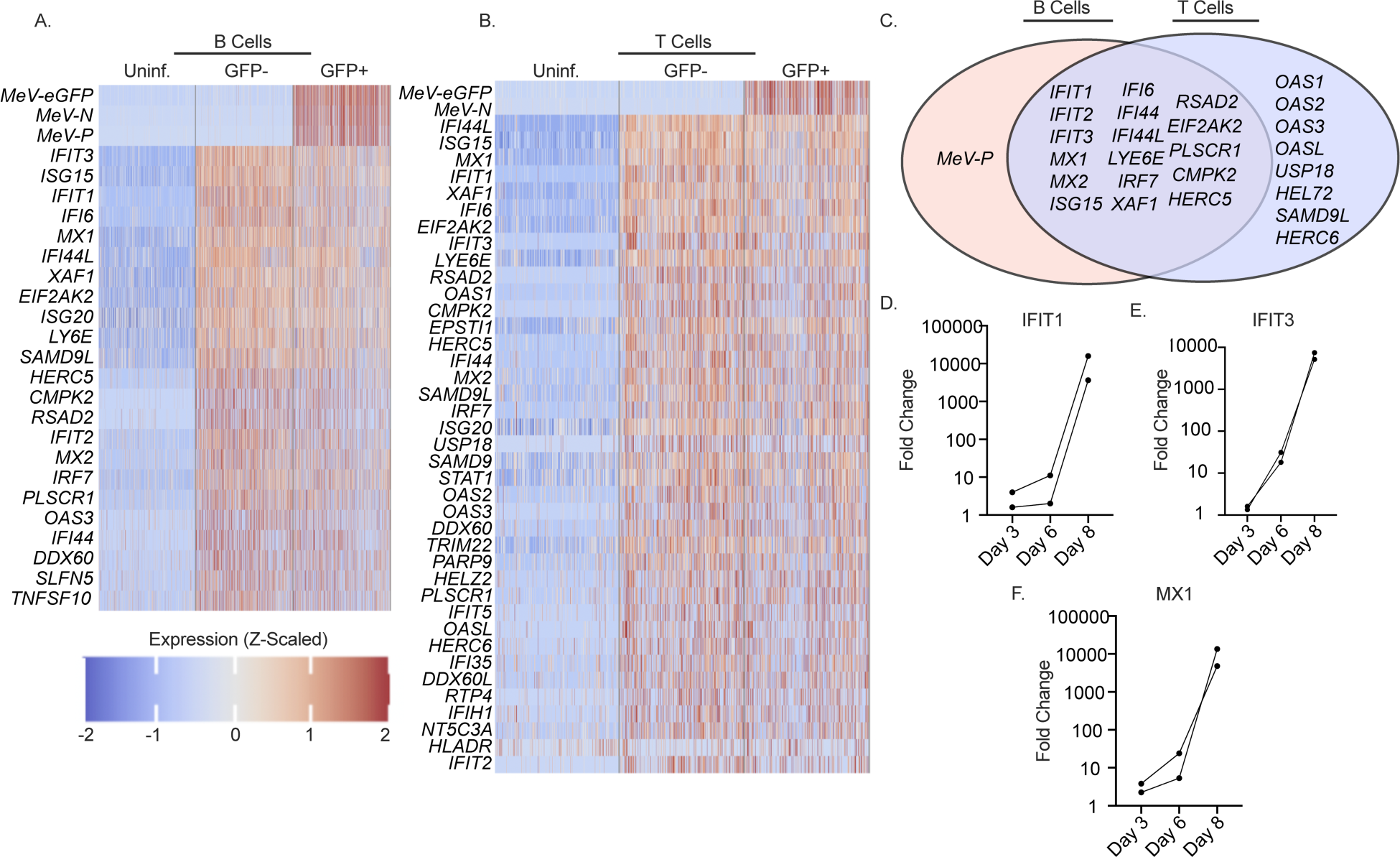
Host response to MeV in lymphoid tissue is dominated by a type I interferon response. All B and T cell clusters from the scRNA-Seq analysis were combined with stochastic downsampling. Differential gene expression analysis was conducted with EdgeR, and expression of the most significant genes is shown for B cells **(A)** and T cells **(B)**. Statistical thresholds for significant differential gene expression were set at Benjamini-Hochberg adjusted *p* value <0.0001 and absolute LogFC>1.58. Genes that were significant in either the infected:uninfected or the bystander:uninfected comparison were plotted. Genes significantly induced during infection were compared between B and T cells **(C)**. MeV-infected tonsil explants were collected for RNA extraction and analysis by qRT-PCR. RNA was examined for the expression of *IFIT1* **(D)**, *IFIT3* **(E)**, and *MX1* **(F)**. The fold change in expression levels relative to uninfected controls is shown, with the expression of each ISG normalized to the expression of *GAPDH* (ΔΔCT method).

### MeV induces an interferon-driven response in B cells at the protein level

Since we identified a potent interferon signature in response to MeV at the transcriptional level, we next asked whether the corresponding proteins were expressed. As B cells were preferential targets for MeV, we utilized Raji cells where we could carefully define the protein level response to infection in a uniform cellular population. Raji cells were infected with MeV at an MOI of 0.1 for

72 hours, and infection was confirmed through GFP expression (**Figure 5A**). Infected and uninfected Raji cells were lysed, trypsin digested and analyzed by quantitative mass spectrometry. Protein abundance was quantified in each sample relative to uninfected samples. We next conducted differential expression analysis to quantify altered protein expression during MeV infection as compared to the uninfected condition. As shown in **Figure 5B**, MeV-infected B cells have higher expression of ISGs, consistent with our scRNA-seq data.

**Figure 5:**
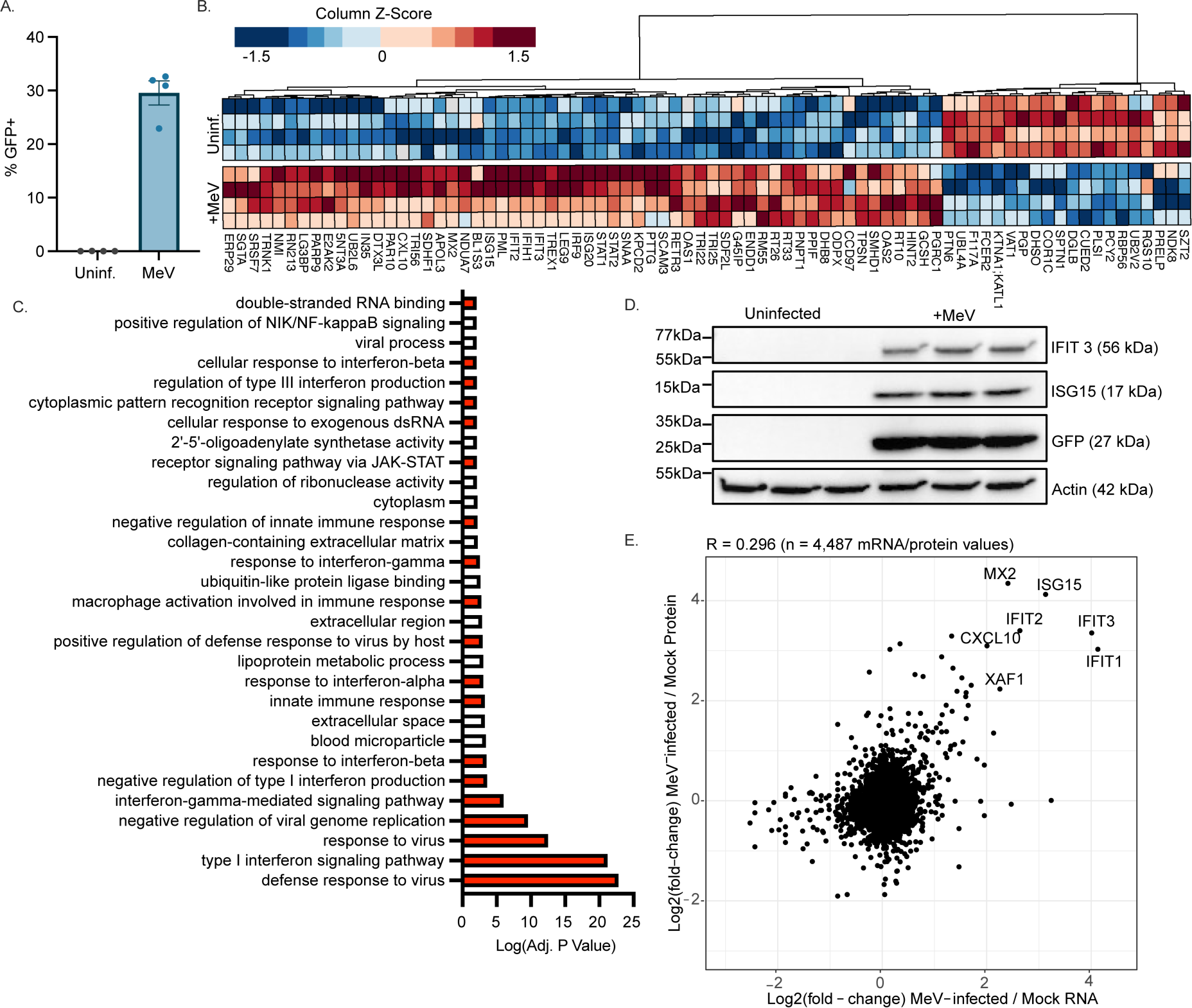
Proteins involved in a type I Interferon response are potently upregulated in response to MeV. Raji-DCSIGNR cells were infected with MeV-GFP (MOI=0.1) or left uninfected for 72 hours before processing for protein abundance mass spectrometry (n=4). Infection was confirmed by quantifying GFP expression by flow cytometry **(A)**. The most significantly dysregulated proteins (columns) for each sample (rows) were visualized, with clustering based on protein expression **(B)**. GO analysis was conducted, and the most significant functional terms were visualized. Terms were given a positive value if the term was upregulated during infection or a negative value if downregulated. Red bars indicate involvement in antiviral responses **(C)**. Validation of IFIT3 and ISG15 upregulation in infected Raji cells was conducted by western blot **(D).** Correlation of the transcriptome (pseudobulked B cell supercluster from Figure 4) and proteome (Raji cells, Figure 5) was conducted. Values that had a log2FC>2 in both the proteome and transcriptome were labeled **(E)**. For the box and whisker plot, the mean with SEM is shown.

To functionally annotate the significantly dysregulated proteins, we next conducted a gene ontology (GO) analysis. Significant GO terms are shown in **Figure 5C**, where the directionality of the response is artificially shown based on a positive (upregulated) or negative (downregulated) transformation of the adjusted p-value for that term. We found that the most significantly upregulated pathways in MeV-infected cells were involved in antiviral signaling or interferon biology (colored in red; **Figure 5C**). To further validate these results, we selected IFIT3 and ISG15, two of the most highly upregulated proteins identified in the proteomics data to examine by western blot. Like the GFP signal, we found that both IFIT3 and ISG15 were expressed only in MeV-infected Raji cells (**Figure 5D**). Given that the transcriptomic analysis was conducted in primary human lymphoid tissue, while the proteomic analysis was conducted in a B cell line, we wanted to identify the significantly upregulated hits that were identified in both systems. To do this, we compared the MeV transcriptional signature among the pseudobulked B cell cluster (containing stochastic sampling of each of the B cell clusters) from lymphoid explants to the significant proteins identified in the proteome of infected Raji cells. As shown in **Figure 5E**, the most highly upregulated hits from this correlation analysis were ISGs, dominated by *MX2*, *ISG15*, *IFIT1*, *IFIT2*, *IFIT3*, *CXCL10* and *XAF1*. To further characterize this host response, we also compared the transcriptome and proteome of infected Raji cells. Raji cells were infected as in the MS experiment but collected for bulk RNA sequencing. As shown in **Figure S7**, we identified a conserved set of interferon stimulated genes that were robustly and significantly upregulated at both the RNA and protein level. In addition, we noted that the expression of the FC-epsilon receptor (*FCER2*) was significantly downregulated. Taken together, our integrated approach of assessing the host response to infection at both the transcriptional and protein level reveals a potently induced interferon signature that is conserved between distinct infection systems.

## DISCUSSION

Measles virus pathogenesis is dependent upon early replication within the draining lymph node, yet our understanding of how infection proceeds in this organ and its link to disease outcomes is incomplete (48). In this study, we sought to model MeV infection in primary human lymphoid tissue explants and comprehensively characterize the immunological events that occur following MeV infection. scRNA-Seq analysis of infected cells was made possible by the high infection rates achieved in this system coupled with a GFP-expressing pathogenic strain of MeV. Our study also contributes an analysis of the MeV-induced proteome within infected B cells. Our unbiased approaches suggest that MeV has a remarkably wide lymphoid tropism, as we found MeV transcripts in most of the 29 cell types identified within the lymphoid tissue explants. Our data confirm previous findings of lymphocyte susceptibility and demonstrate a strong IFN signature associated with infection at both the RNA and protein levels.

Previous studies on MeV pathogenesis have identified a bias in infection towards B cells and away from T cells, with susceptibility differences driven by the expression of CD150 (18, 20, 27, 34, 40, 48, 49). Our data are consistent with this notion, as we found that B cells were the largest targets of infection, having heightened susceptibility and trending higher CD150 expression compared to T cells. Our analysis also extends these findings, revealing that while all B cells are highly susceptible to infection, accounting for the majority of infected cells, there were no observed differences in susceptibility based on B cell phenotype. Since CD150 expression was generally high among these B cell subsets, we interpret these findings to indicate that a baseline level of CD150 expression is sufficient to confer susceptibility, but differences beyond this threshold do not alter susceptibility. Of note, we found that germinal center B cells (GCBs) were by far the largest population of B cells found in this tissue, and thus, also comprised the greatest number of MeV-infected cells. These findings may suggest that immunological amnesia may extend beyond existing immunological to hamper future germinal center responses.

An interesting observation that we uncovered was that all cell types that we identified in the tonsil cultures were found to have some level of MeV transcripts. Several cell types had extremely low levels of MeV transcripts, including stromal and epithelial cells. While we took many measures to ensure that these were bona fide infected cells (such as dead cell removal, flow sorting for GFP, and a careful hashing strategy), we cannot eliminate the possibility that these cells are a byproduct of exosomes or ambient viral RNA co-encapsulated with the GFP^-^ populations rather than truly infected cells. Our findings in larger populations, such as subsets of CD4 and CD8 cells, were recapitulated with our flow cytometry approach, however, future work should assess the possibility of MeV infection in rare tonsillar populations, such as stroma and epithelium.

Many early studies on MeV pathogenesis focused on infection of T cells within secondary lymphoid tissue (21, 34, 37, 43, 49, 50). Indeed, immunological amnesia was originally described as a T cell phenotype, where children who had previously tested positive for a hypersensitivity test to tuberculin antigen began to test negative following MeV infection (51). Subsequent work in thymus and macaques revealed that MeV preferentially infects and depletes memory T cells over naive T cell subsets, consistent with CD150 expression (32, 41). Our approach of assessing the relative susceptibility of both T cells broadly, as well as within individual subsets both confirmed and extended these findings. We found that antigen experience (CD45RA negativity) influenced susceptibility to infection, while CXCR5 expression, used here as a proxy for localization within the lymphoid explants, as well as CD150 expression, did not. Future work assessing the susceptibility of these antigen-experienced subsets should focus on directly testing if factors other than CD150 expression, such as spatial localization, promote susceptibility. Indeed, one parameter that may be interesting to evaluate would be the extent to which directed cell migration occurs within the tissue, and if infection influences immune cell trafficking.

Previous groups have shown that MeV does not induce a potent interferon (IFN) response, as the viral V and C proteins can inhibit the induction of IFN (52–60). However, some groups have observed the opposite, where MeV induces potent IFN expression (46, 61, 62). In general, this discrepancy has been attributed to the presence of defective interfering (DI) RNAs, which can be enriched as a byproduct of in vitro replication (52, 59, 63, 64). While these would not be captured by our scRNA-Seq modality, we can conclude that the presence of viral V transcripts at day 8 was not sufficient to shut down the IFN response, whether induced by DI RNAs or viral replication. One hypothesis that might explain how MeV replicates in the presence of an IFN response would be that the IFN response is induced to the benefit of MeV, not the detriment. The idea that viruses may utilize IFN responses to promote infection has recently been demonstrated for influenza virus, whereby the virus utilizes the host ISG *IFIT2* to enhance the translational efficiency of viral RNAs (65). Alternatively, the addition of GFP into the viral genome may be indirectly involved as placement of GFP in the first transcriptional unit of our MeV-GFP may decrease the relative amounts of P-derived V and C proteins that antagonize type-I IFN responses. MeV-C is known to reduce the production of DIs by enhancing the processivity of the viral polymerase (66) with C-deficient MeV generating ∼10-fold more DIs than the parental virus (64). Future studies should assess the impact of this transcriptional shift on DI production as well as the downstream ability to antagonize the type-I IFN response.

One major limitation of our study is that we do not know the impact of MeV infection disease outcomes, such as immunological amnesia. Our results suggest that MeV infection of GBCs may impact the germinal center responses, an outcome that would amplify the impact of MeV on immunological amnesia. A second limitation of our study is that the transcriptomic and proteomic analyses were conducted in entirely different systems due to the heterogeneous nature of the lymphoid tissue explants. In the absence of single-cell proteomics, we limited our approach to a correlative analysis between the two methodologies and systems. Therefore, we have high confidence that these molecules are indeed a conserved response to MeV infection. A third limitation in our analysis is that we are unable to differentiate between cell death by MeV versus cell susceptibility to infection. While our data suggest that the broad susceptibility of T and B cells is positively associated with the expression of the entry receptor CD150, this does not exclude the possibility that some cell subpopulations have a greater capacity to survive while infected with MeV. Indeed, it has previously been established that MeV is capable of depleting CD150^+^ cells in tonsil explants (43). However, given the high similarity in the frequency of bystander cells and uninfected cells across all cell populations identified, we can presume that an enhanced frequency among GFP^+^ cells is indicative of enhanced susceptibility to infection. Understanding the impact of MeV on cell death and proliferation may prove critical to understanding the complete pathology of measles disease. Another limitation of our study is that we do not know the impact of the GFP produced by the MeV-GFP on the induction of the innate immune response in tonsil tissues and/or cell lines. The use of the GFP-expressing MeV enabled us to distinguish between infected and bystander cells in our culture system. This was unavoidable to sort and conduct scRNAseq for this study. However, a study utilizing this MeV-GFP showed that this strain is fully pathogenic in macaques, suggesting that the introduction of GFP, and its possible ISG induction, does not affect viral pathogenesis (67).

Our findings here represent a thorough analysis of the immunological events following MeV infection of human lymphoid tissue explants. The finding that MeV has a broad tropism within B cell, T cell, myeloid, and non-hematopoietic compartments may unlock new aspects of viral pathogenesis in humans. While we do not know the role of each of these cell types in the collective immunological response to infection, future studies should investigate how these cell types shape the progression of measles disease. Our findings also represent a model system for the testing of MeV antivirals, for which there are no current intervention strategies. One possibility would be that by targeting specific aspects of the induced IFN response, MeV pathogenesis could be ameliorated.

In toto, we present a thorough kinetic examination of the process of MeV infection in human lymphoid explants, confirming previous groups’ findings and broadening our understanding of the key players in MeV infection within its natural target organ architecture. Further, our integrated transcriptional and proteomic approach in this model establishes tonsil explants as a unique platform for the identification of host factors important for MeV replication and screening of targeted antivirals. Future work in this model should focus on understanding how MeV replicates in the face of this potently induced interferon response to identify junctions at which viral replication can be inhibited.

## MATERIALS AND METHODS

### Sex as a biological variable

Our study received human tonsil tissue from both males and females. We did not observe any clear difference in MeV replication, so these data were analyzed together.

### Cells and plasmids

Vero-hCD150 cells were provided by Dr. Yanagi at Kyushu University and maintained in DMEM with 10% FBS (Biowest). RAJI-DCSIGNR cells were gifted by Ted Pierson (NIH/VRC, Bethesda USA) and cultured in RPMI with 10% FBS (68). The genome coding plasmid for MeV (p(+) MV323-AcGFP) was kindly gifted from Dr. Makoto Takeda (University of Tokyo, Tokyo Japan) (47). The MeV genome sequence was transferred into a pEMC vector, adding an optimal T7 promoter, a hammerhead ribozyme, and an eGFP transcriptional unit at the 3’ end of the genome (pEMC-IC323-eGFP) as previously described (47).

### MeV rescue and amplification

MeV (IC323-eGFP) rescue was performed in BSR-T7 cells, seeded in 6-well format. Upon confluency, pEMC-IC323eGFP (5μg), T7-MeV-N (1.2μg), T7-MeV-P (1.2μg), T7-MeV-L (0.4μg), a plasmid encoding a codon-optimized T7 polymerase (3μg), PLUS reagent (5.8μL, Invitrogen), and Lipofectamine LTX (9.3μL, Invitrogen) were combined in Opti-MEM (200μl; Invitrogen). After a 30 min incubation at RT, the transfection mixture was added dropwise onto cells and incubated for 24hrs at 37°C. Following, rescued virus was amplified once on Vero-hCD150 cells for 72hrs to generate a P1 virus, in infection media (made in DMEM+2% FBS). This virus was then titered (see plaque assay method below) and used at an MOI = 0.01 on Vero-hCD150 cells to generate a P2 virus (amplified as above). Supernatants were collected, clarified of cell debris, ultracentrifuged through a 20% sucrose gradient at 24,000 RPM for 3hrs, reconstituted in fresh infection media, and frozen at −80°C.

### MeV quantification by plaque assay

Vero-hCD150 were plated in 12 well format until ∼90-95% confluence. 10-fold dilutions of samples (made in DMEM +2% FBS) were applied to these monolayers in a total volume of 250µL, and infections were allowed to incubate for 2hrs at 37°C. Viral inoculum was replaced with 500µL/well of methylcellulose (in DMEM + 2% FBS + 7.5% NaHCO_3_). At 72hrs, wells were imaged for GFP^+^ plaques on the Celigo S platform.

### Processing and infection of human lymphoid tissue

Human tonsils from routine tonsillectomies performed at the Mount Sinai Hospital and the New York Eye and Ear Infirmary of Mount Sinai were collected under IRB-approved protocols within a few hours after surgery. Tonsils were cut into 2mm^3^ blocks, and 9 tissue blocks per well were placed on top of collagen gelfoams (Cardinal Health) in a 6-well plate as previously described, utilizing three wells per condition (a total of 27 blocks per experimental sample) (43). In all experiments, triplicate wells were harvested as a single sample to reduce variability (44). After overnight incubation, individual tissue blocks were individually inoculated with 5μl containing 1,666 PFU MeV-GFP (for a final concentration of 5,000 PFU/ml) or left uninfected. Media was collected and replaced at days 3, 6, and 8 post-infection. Tonsil donors consisted of three male and eight female donors. The median age of donors was 23 years old, with a range of 4 to 54 years old. The reasons for tonsillectomy included sleep apnea, breathing disorders, and chronic tonsillitis.

### Visualization of MeV-infected cells in tonsillar explants by in situ hybridization

Tissues were fixed in 10% neutral buffered formalin and paraffin-embedded. In situ hybridization using RNAscope® (ACDBio) was performed on 5μm sections to detect RNA encoding GFP. Deparaffinization was performed by baking slides at 55°C for 20 min. Slides were washed twice with xylene, twice in 100% ethanol, and were dried for 5 min at 60°C. Slides were then incubated with hydrogen peroxide for 10 min at RT and were subsequently washed in diH2O. Slides were placed in Target Retrieval solution at 100°C for 15 min, washed with water, and transferred into 100% ethanol for 3min, before drying. Sections were treated with RNAscope® Protease Plus and fluorescence in situ hybridization was subsequently performed according to the manufacturer’s protocol (ACD# 323110) with RNAscope® Probe EGFP (ACD #400281; binds eGFP RNA) as previously described (69). Slides were then mounted with Vectashield hard-set mounting medium with DAPI (Vector Laboratories) and analyzed using an AxioImager Z2 microscope (Zeiss) and Zen 2012 software (Zeiss).

### Generating single cell suspensions from tonsil histocultures

Single cell suspensions were generated by dissociating tissue (merged from the three technical triplicate wells) using Collagenase IV (Worthington Biochemical) incubated for 30 min at 37°C with gentle shaking as previously described (44). Samples were homogenized with mortar and pestle before filtration over a 100µm cell filter and washed once with cold PBS before downstream application.

### Single cell RNA sequencing (scRNA-Seq)

Samples for scRNA-Seq were pooled for multiplex processing and analysis with a cell hashing antibody strategy (70). Hash antibodies were generated by conjugating IDT synthesized oligos (barcode sequences from 10x Genomics Chromium index SI-GA-F11; HBC21-29 for hash #1-8) to antibodies utilizing Thunder-Link® PLUS oligo Antibody Conjugation Kit. Single cell suspensions were generated from one donor-matched infected and uninfected culture at day 8 and dead cells were depleted from samples using the EasySep™ Dead Cell Removal (Annexin V) Kit (Stemcell Technologies, #17899). Cells were blocked with Human TruStain FcX^TM^ (Biolegend, #422302). Cells from the uninfected tonsil were split into 4 hashing groups (hash #1-4), and cells from the infected tonsil were split into two infected hashing groups (hash #5, 6) and two bystander hashing groups (hash #7, 8). Samples were stained with corresponding hashing antibodies for 30 min at 4°C and washed 3x in FACS buffer (PBS + 1mM EDTA + 2% BSA). Cell suspensions were filtered over a 70µm filter, stained with propidium iodide (PI) for viability, and sorted as follows: live/GFP^-^ cells from the uninfected condition, live/GFP^+^ cells from the infected condition, and live/GFP^-^ cells from the bystander condition on a BD FACS Aria III. Sorted cells were counted, and 10,000 uninfected cells, 5,000 infected cells, and 5,000 bystander cells were pooled and processed for scRNA-Seq on the 10x Genomics Chromium platform, utilizing the 10x 3’ v3 kit. A scRNA-Seq library was generated as per the manufacturer’s protocol and sequenced on an Illumina NextSeq500 instrument. A corresponding library of barcoded hash antibody oligonucleotides was indexed with a standard Illumina D701 index and sequenced as above.

### Processing of scRNA-Seq data

Raw sequencing data output (BCL files) was converted to fastq files with CellRanger mkfastq v3.0.2 (10X Genomics). Per cell gene count and hashtag antibody count matrices were generated with CellRanger count v3.0.2 (10X Genomics), using a human genome reference (GRCh38, Ensembl v96 transcript annotations) appended with the MeV-eGFP reference and corresponding transcript annotations (MeV-IC323-eGFP, Genbank: MW401770). Data were read into the R statistical framework (v4.0.3) for additional analysis with Seurat(71, 72) (v4.0.1). Hashtag antibody data were center log ratio normalized by feature, and individual samples were demultiplexed with the Seurat HTODemux function with the positive.quantile parameter set to 0.99.

### Quality control and filtering of scRNA-Seq

Data exploration and HTODemux classifications were used to set QC thresholds on per cell transcript unique molecular identifiers (UMI) counts, detected gene counts, and the percentage of detected mitochondrial transcripts. Cells with fewer than 2,500 transcript UMIs, fewer than 800 detected genes, and greater than 15% mitochondrial transcripts were excluded from downstream analyses. After filtering, these data included 5,737 cells in the uninfected group, 2,736 cells in the infected group, and 2,944 in the bystander group.

### scRNA-Seq data analysis

Datasets were normalized with SCTransform (73), with the per-cell mitochondrial transcript percentage included as a regression variable. Data from all groups were integrated in Seurat using 3,000 anchor features; MeV genes were excluded from all integration and clustering steps to avoid group-specific artifacts. Dimensionality reduction was performed by principal component analysis on integrated data, and the first 20 components were selected for graph-based clustering by smart local moving algorithm (74) at a resolution of 1.4 (determined by Clustering Tree assessment (75)).

General cell types were annotated by SingleR (76) from human immune cell reference data (77). Clusters were assigned to one of each major cell group: T/NK, B, plasma, myeloid, stromal, and epithelial. Those major cell groups with multiple component clusters were subset and re-analyzed (normalization, principal components analysis dimensionality reduction, and clustering at ClusterTree determined optimal resolution) for further annotation. For each major cell group subset analysis, “marker genes” distinguishing component clusters were identified with the FindAllMarkers (on the uninfected group) and/or FindConservedMarkers (on all groups) functions. Intergroup differential gene expression analysis was performed with edgeR (78, 79) (v3.32.1), including modifications of scRNA-Seq data (80). The edgeR linear model incorporated factors for cellular gene detection rate (to account for scRNA-Seq “dropout”) and experimental group and included only those genes detected in at least 20% of cells in any contrast condition. Statistical thresholds were set at Benjamini-Hochberg adjusted *p*-value less than 0.0001 and absolute log fold-change greater than 1.58 for differential expression.

### Immunophenotyping by flow cytometry

Cells were stained with the Zombie Red fixable viability kit (Biolegend, #423109) for 10min at RT, washed once with FACS buffer, and then blocked with Human TruStain FcX^TM^ (Biolegend, #422302). Samples were incubated for 30min on ice with a cocktail of antibodies against: **(B Cell Panel)** CD150 (Biolegend; Clone: A127d4; PE), CD38 (eBioscience; Clone: HB7; PE-Cy7), CD27 (Biolegend; Clone: O323; APC), CD45 (BD Horizon; Clone: HI30; BV605), CD19 (Biolegend; Clone: HIB19; BV750) and CD3 (Biolegend; Clone: OKT3; BV785); **(T Cell Panel)** CD4 (eBioscience; Clone: OKT4; PerCP-Cy5.5), CD150 (Biolegend; Clone: A127d4; PE), CD45RA (Biolegend; Clone: HI100; AlexaFluor-700), CXCR5 (Biolegend; Clone: J252D4; BV421), CD8 (Biolegend; Clone: RPA-T8; BV570), CD45 (BD Horizon; Clone: HI30; BV605), CD19 (Biolegend; Clone: HIB19; BV750) and CD3 (Biolegend; Clone: OKT3; BV785). The antibody cocktail was supplemented with Brilliant Stain buffer (BD Horizon^TM,^ #563794). All antibodies were used at a concentration of 1μg/ml, except for CD27 (4μg/ml). Cells were washed 3 times with FACS buffer before fixation with BD Cytofix^TM^ (BD Biosciences; #554655). Single color controls were generated on UltraComp eBeads™ (Invitrogen™, #01-2222-42), except for GFP and Live/Dead controls, which were generated using cells. All samples were analyzed on an Aurora Cytek®, and unmixed samples were analyzed in FlowJo v10.8.1.

### qRT-PCR

Resuspended single tonsil cell suspensions were placed in 1ml of TriZol, and RNA was isolated using Direct-zol RNA MiniPrep Plus kit (Zymo). 1µg of RNA was reverse transcribed with random hexamer primers (Applied Biosystems). 1µl of cDNA was utilized per reaction, and primer/probes for *IFIT1* (HS03027069_S1), *IFIT3* (HS01922752_S1), and *MX1* (Hs00895608_m) were utilized to amplify ISG transcripts. Fold induction was calculated using the ΔΔCT method by normalizing expression to *GAPDH* expression (NC_000012.11).

### Sample preparation for Mass Spectrometry

Uninfected or MeV-infected Raji-DCSIGNR cells were lysed in an 8M urea lysis buffer (with 100 mM ammonium bicarbonate, 150 mM NaCl, and 1x protease/phosphatase inhibitor cocktail HALT (Thermo Fisher Scientific)). Lysates were sonicated, and protein concentrations were quantified by micro-BCA assay (Thermo Fisher Scientific). 50μg of protein for each sample were treated with Tris-(2-carboxyethyl)phosphine (TCEP) at a 4mM final concentration and incubated for 30 min at RT. Iodoacetamide (IAA) was added to a 10mM final concentration and samples were incubated for 30 min at RT. Free IAA was quenched with the addition of Dithiothreitol (DTT) at a 10mM final concentration for 30min. Samples were diluted with 5 sample volumes of 100 mM ammonium bicarbonate. Lysates were next digested with Trypsin Gold (Promega) at a 1:100 (enzyme: protein) ratio and lysates were rotated for 16hrs at RT. Trypsin activity was quenched by adding 10% v/v trifluoroacetic acid (TFA) to a final concentration of 0.1%. Samples were desalted on BioPure™ SPN MIDI C18 Spin columns. Samples were eluted from these columns with 200µL 40% ACN/0.1% TFA, dried by vacuum centrifugation, and stored at −80°C.

### Protein abundance Mass Spectrometry

Samples were analyzed on an Orbitrap Eclipse mass spectrometry system (Thermo Fisher Scientific) equipped with an Easy nLC 1200 ultra-high pressure liquid chromatography system (Thermo Fisher Scientific) interfaced via a Nanospray Flex nanoelectrospray source. Immediately before spectrometry, lyophilized samples were resuspended in 0.1% formic acid. Samples were injected on a C18 reverse phase column (30 cm x 75 μm (ID)) packed with ReprosilPur 1.9 μm particles). Mobile phase A consisted of 0.1% FA, and mobile phase B consisted of 0.1% FA/80% ACN. Peptides were separated by an organic gradient from 5% to 35% mobile phase B over 120 min followed by an increase to 100% B over 10 min at a flow rate of 300 nL/min. Analytical columns were equilibrated with 3 μL of mobile phase A. To build a spectral library, samples from each set of biological replicates were pooled and acquired in a data-dependent manner. Data-dependent analysis (DDA) was performed by acquiring a full scan over a m/z range of 375-1025 in the Orbitrap at 120,000 resolution(@200 m/z) with a normalized AGC target of 100%, an RF lens setting of 30%, and an instrument-controlled ion injection time. Dynamic exclusion was set to 30 seconds, with a 10ppm exclusion width setting. Peptides with charge states 2-6 were selected for MS/MS interrogation using higher energy collisional dissociation (HCD) with a normalized HCD collision energy of 28%, with three seconds of MS/MS scans per cycle. Data-independent analysis (DIA) was performed on all individual samples. An MS scan was performed at 60,000 resolution (@200m/z) over a scan range of 390-1010 m/z, an instrument-controlled AGC target, an RF lens setting of 30%, and an instrument-controlled maximum injection time, followed by DIA scans using 8 m/z isolation windows over 400-1000 m/z at a normalized HCD collision energy of 28%.

### MS Data analysis

Peptides/Proteins were first identified with Spectronaut (81). False discovery rates (FDR) were estimated using a decoy database strategy. All data were filtered to achieve an FDR of 0.01 for peptide-spectrum matches, peptide identifications, and protein identifications. Search parameters included a fixed modification for carbamidomethyl cysteine and variable modifications for N-terminal protein acetylation and methionine oxidation. All other search parameters were defaults for the respective algorithms. Analysis of protein expression utilized the MSstats statistical package in R. Output data from Spectronaut was annotated based on a publicly available *Homo sapiens* proteome (Proteome ID: UP000005640) and the reference sequence for IC323-eGFP (Genbank: MW401770.1). Technical and biological replicates were integrated to estimate log2 fold changes, p values, and adjusted p values. All data were normalized by equalizing median intensities, the summary method was Tukey’s median polish, and the maximum quantile for deciding censored missing values was 0.999. Significantly dysregulated proteins were defined as those that had a fold change value >2 or <-2, with a p-value of <0.05. The mass indices for the most significantly dysregulated proteins were transformed with the Quantile function in R and then visualized using the pheatmap package.

### Gene Ontology (GO) analysis

Gene Ontology (GO) analysis was conducted in R by quantifying the number of significantly upregulated or downregulated proteins for each GO term identified from the gene ontology resource, downloaded on 2.18.21. Significance was determined by a hypergeometric test. Only terms with >2 proteins detected in the dataset were included in this analysis. This approach was similarly taken to identify significantly downregulated terms. GO terms were graphed in GraphPad PRISM based on their p-value, and terms related to the antiviral response were colored in red.

### Western blot analysis of RAJI cell lysates

Raji cells were infected with MeV at an MOI= 0.1 for 72 hours and compared to uninfected controls (n=3). Cells were lysed as described above, and 10ug samples of whole cell lysate were mixed 1:1 with Lamelli Buffer (containing b-mercaptoethanol) and were heated at 95°C for 10min. Samples were then electrophoresed on a 4-20% gradient SDS-PAGE gel (BioRad) and transferred onto a methanol-activated polyvinylidene difluoride (PVDF) membrane (Bio-Rad). The membrane was blocked with 5% Milk in PBST (0.1% Tween-20) for 1hr. The following antibody staining protocols were run sequentially: 1) anti-IFIT3 (ThermoFisher; Clone: OTI1G1; 1:1000) developed with goat-anti-mouse IgG-HRP (Catalog: G-21040; 1:10,000); 2) anti-ISG15 (Clone: 7H29L24; 1:5000) developed with goat-anti-rabbit IgG-HRP antibody (Thermofisher; Catalog: 65-6120; 1:10,000); and 3) anti-beta Actin (Thermofisher; Clone: 15G5A11/E2; 1:1000) and anti-GFP (Thermofisher; Clone: GF28R; 1:1000) simultaneously, developed with anti-mouse Alexa 647 antibody (Thermofisher; clone A-21235; 1:2000). HRP signals were detected between each incubation with SuperSignal West Pico™ PLUS reagent (Thermofisher; 1:1 luminol/enhancer), and images were acquired on a Chemidoc™ MP. Western images were merged for presentation in FIJI.

### Correlation Analysis of RNA and protein response to infection

Data from scRNAseq and mass spectrometry were further processed in RStudio to correlate RNA and protein levels. All B cells from the scRNAseq dataset were rebulked, and total read counts in this new “B cell” cluster were calculated, normalized to counts per kb million, and log2 transformed. To evaluate the fold change (FC) between mock-treated and MeV-infected samples, the log2 values from the mock condition were subtracted from the MeV condition. A correlation scatter plot was created using ggplot, and protein labels were added only if the log2FC values were greater than 2 in both RNA and protein.

### Bulk-RNA sequencing of infected RAJI cells

RAJI-DCSIGNR cells were infected with MeV as during the preparation of MS samples above. Cells were pelleted and resuspended in 500uL of Trizol. RNA was extracted using the Direct-zol RNA miniprep kit (Zymo Research), and frozen RNA was shipped to Azenta Life Sciences for library preparation and sequencing. ERCC RNA Spike-in Max kit was added to normalize total RNA prior to library preparation following the manufacturer’s protocol (cat # 4456740). RNA sequencing libraries were prepared using the NEBNext Ultra II RNA Library Prep Kit for Illumina. Libraries were quantified with Agilent TapeStation, Qubit 2.0 and by quantitative PCR prior to sequencing on an Illumina NovaSeq XPlus 25B. Samples were sequenced using a standard 2×150bp paired-end configuration. Raw sequence data was converted into fastq files and de-multiplex using Illumina’s bcl2fastq 2.2.0 software. The quality of sequencing was assessed with FastQC, and sequencing reads were aligned to indexed reference genomes using the STAR aligner. Expression matrices were calculated using featureCounts, and data were exported into R for data analysis and visualization. Transcripts where fewer than 10 transcripts were collected across all samples were excluded from further analysis. Data normalization and differential gene expression analysis was conducted using the DESEQ2 package (version 1.42.1).

### Statistics

For scRNA-Seq and mass spectrometry analysis, statistical analysis methodology is detailed in the above methods section. For all comparisons of infection susceptibility over time, significance was determined by two-way ANOVA using the Geisser-Greenhouse correction with Tukey’s multiple comparison test. For all comparisons of CD150 expression among multiple (>2) groups, significance was determined by one-way ANOVA using Friedman’s test with Dunnett’s multiple comparison test. For pairwise comparisons, a non-parametric 2-tailed paired T-test was utilized (Wilcoxon matched-pairs signed rank test). For experiments with three replicates, the median with the 95% confidence intervals is shown instead of p-value. For all other statistical analyses, * indicates *p*<0.05, ** indicates *p*<0.01, *** indicates *p*<0.001 and **** indicates *p*<0.0001

### Study Approval

All tonsil tissues were obtained with informed consent under IRB# 16-01425-CR002 at the New York Eye and Ear Infirmary of Mount Sinai, or the Mount Sinai Hospital under IRB HS#12-0045.

## Data Availability

Large data sets will be made available on NCBI GEO and SRA. BulkSeq (Accession #GSE272426); scRNASeq (Accession #GSE272481). Proteomics data are available on PRIDE. Raw data values for data shown in this manuscript can be accessed in the “Supporting data values” XLS file. Information about human subjects is limited by IRB, however anonymized information will be made available upon request to the corresponding author where possible.

## AUTHOR CONTRIBUTIONS

JAA, BRR, BL and JKL conceptualized the project. JA, ARP, SH, APK, PT, SI, JC, HU, BT, JRJ, BRR, BL, and JKL contributed to the work intellectually. JAA, ARP, SH, ASM, PT, and NI conducted the experiments. SI generated the virus used in this study. BT conducted tonsillectomies included in this work. JAA, ARP, SH, EJD, and BRR conducted data analysis. JAA, ARP, and JKL prepared the manuscript. All authors edited the manuscript. The order of the co-first authors was determined based on contributions to the conceptualization and execution of this work.

## ACKNOWLEDGMENTS

Research was supported by NIH applications R21AI149033 (JKL, BRR, and BL), R01AI071002 (BL), and CDMRP PR192188 (JKL and BL). JAA was supported by an NIH fellowship F31HL149295. ARP and EJD were supported in part by the NIH training grant T32AI007647-23. P.A.T. was funded by CIHR-IRSC: 0041001056. The authors thank Dr. Rachel Brody and colleagues at the ISMMS Biorepository and Pathology CoRE, the ISMMS Human Immune Monitoring Core (HIMC), and Dr. Emilia Bagiella, for her expertise and guidance on the statistical methodology.

---

**Figure S1 (related to Figure 2):**
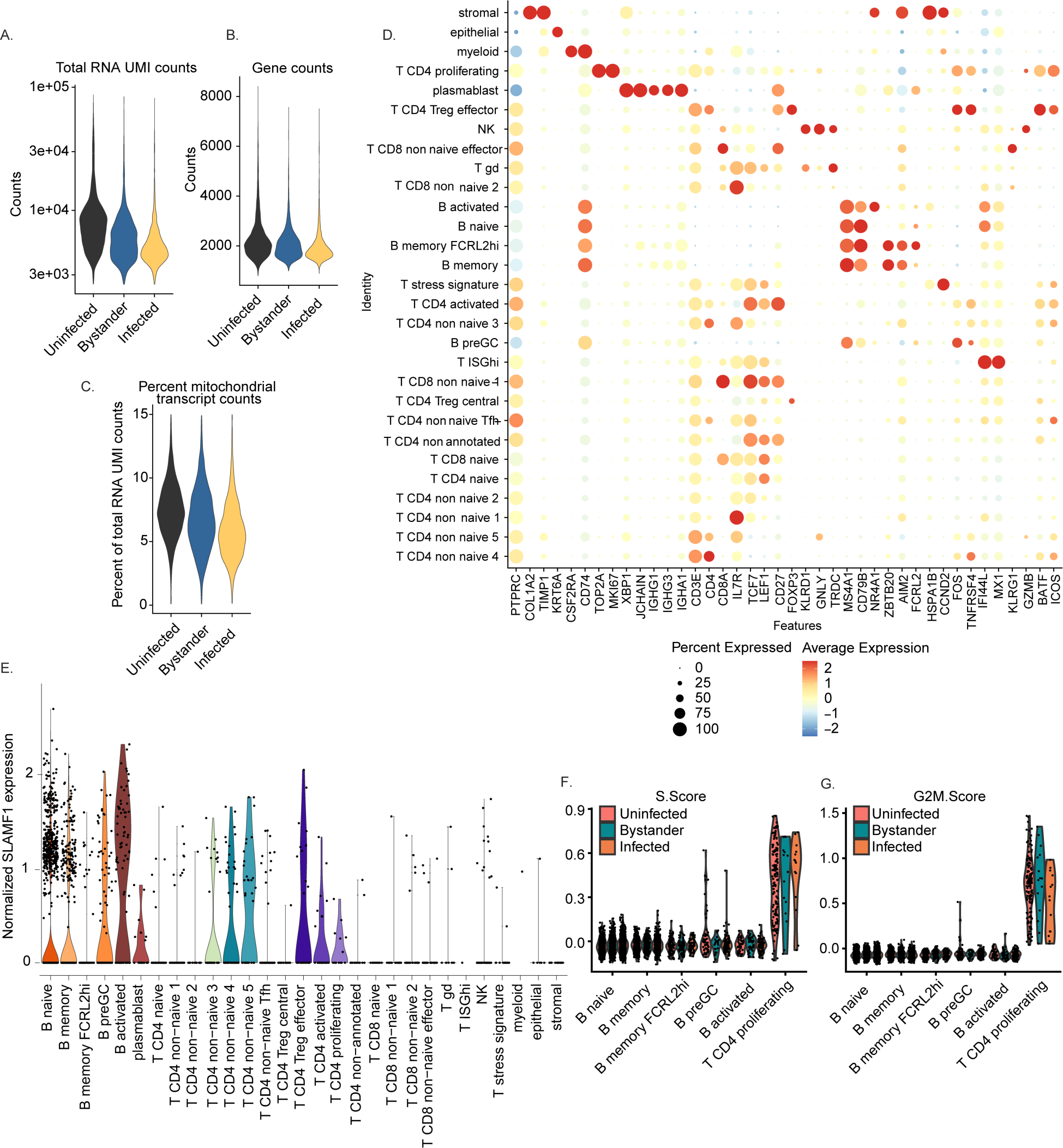
Cluster annotation strategy for scRNA-Seq. Total counts of unique molecular identifiers (UMI; **A**), individual genes **(B)**, and mitochondrial transcripts **(C)** were quantified for quality control and filtering. Dot plot showing the expression of cluster-defining features across each identified cell cluster. Average expression (Z scaled) is shown for each feature by color, while the percentage of cells in each cluster expressing that feature is shown by the size of the dot **(D)**. Normalized SLAMF1 expression across all cell types was determined **(E)**. Pathway scores for S phase **(F)** and G2M **(G)** were calculated and compared across B cell subsets and proliferating CD4 T cells.

**Figure S2 (Related to Figure 3):**
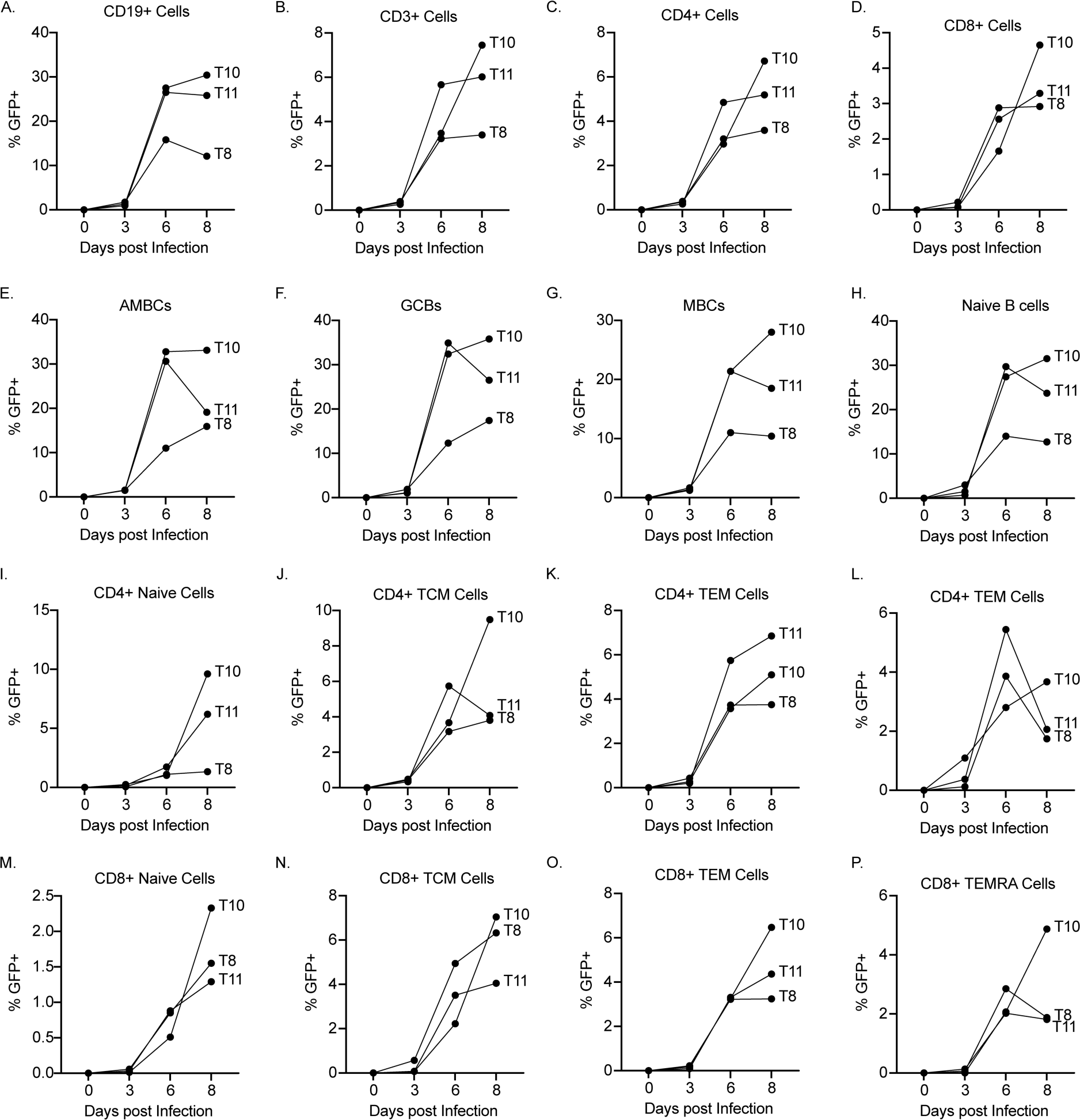
Quantification of infection kinetics in different cell populations. Secondary analysis of donors from Figure 3. The percentage of GFP^+^ cells within each cell type identified in the explants is quantified for B and T cells **(A-B)**, CD4^+^ and CD8^+^ cells **(C-D)**, B cell subsets **(E-H)**, CD4^+^ T cell subsets **(I-L)** and CD8^+^ T cell subsets **(M-P)**. Each dot/line represents a single donor.

**Figure S3 (Related to Figure 3).**
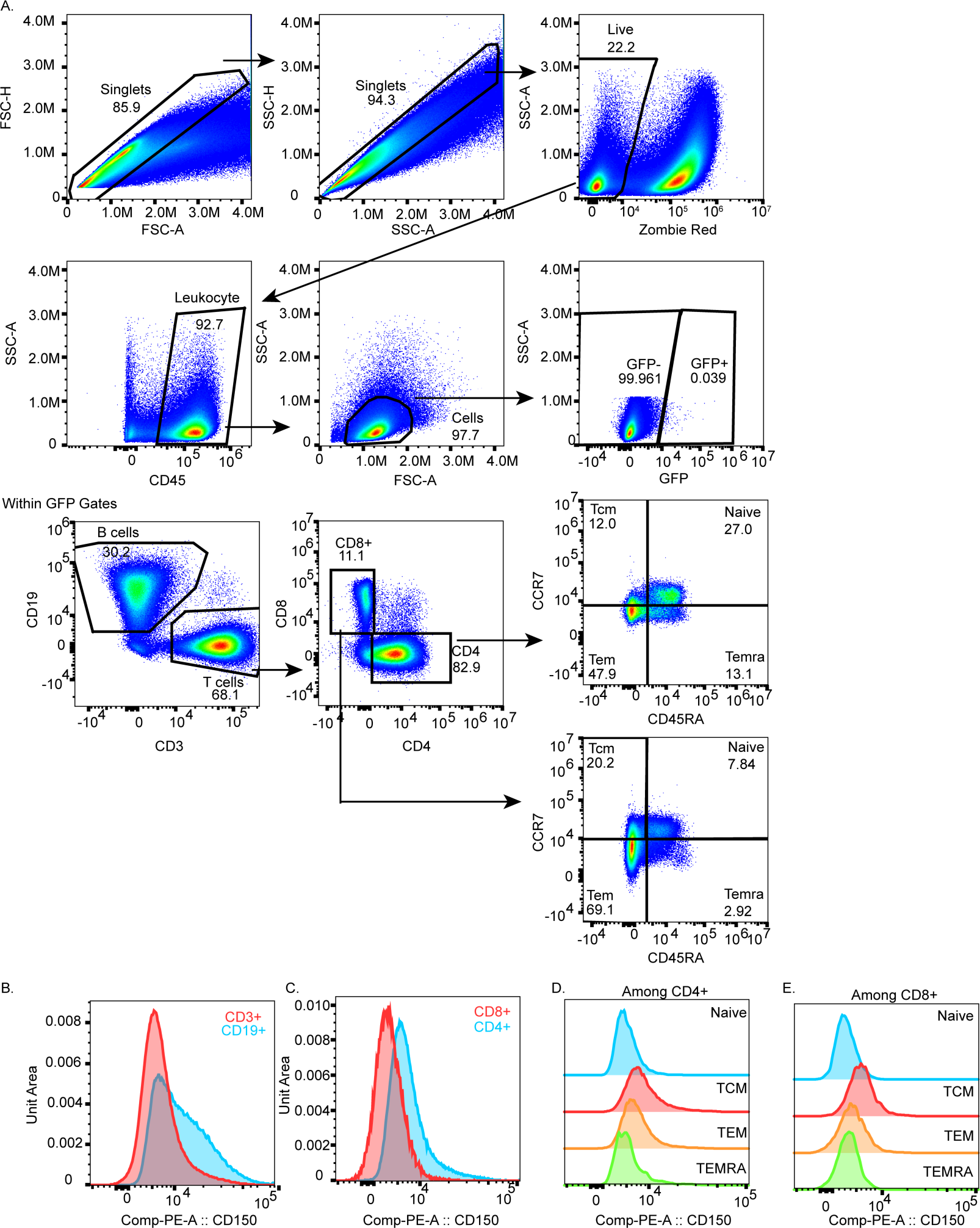
Gating schemata for flow cytometry-based immunophenotyping experiments. Representative flow plots are shown for one uninfected donor. Single cells are selected, followed by dead cell exclusion, gating upon CD45^+^ cells, cell gating, and then binning cells based on GFP status **(A)**. Within GFP^+^ or GFP-gates, cells are further phenotyped based on CD3/CD19 (B vs T cells), CD4 vs CD8 lineages within the CD3^+^ gate, and then memory subsets within these lineages. Histograms demonstrating CD150 expression are compared between CD3^+^ and CD19^+^ cells **(B)**, between CD4^+^ and CD8^+^ cells **(C)**, or between CD4^+^ **(D)** or CD8^+^ **(E)** memory subsets for one representative donor in the day 6 uninfected condition.

**Figure S4 (related to Figure 3):**
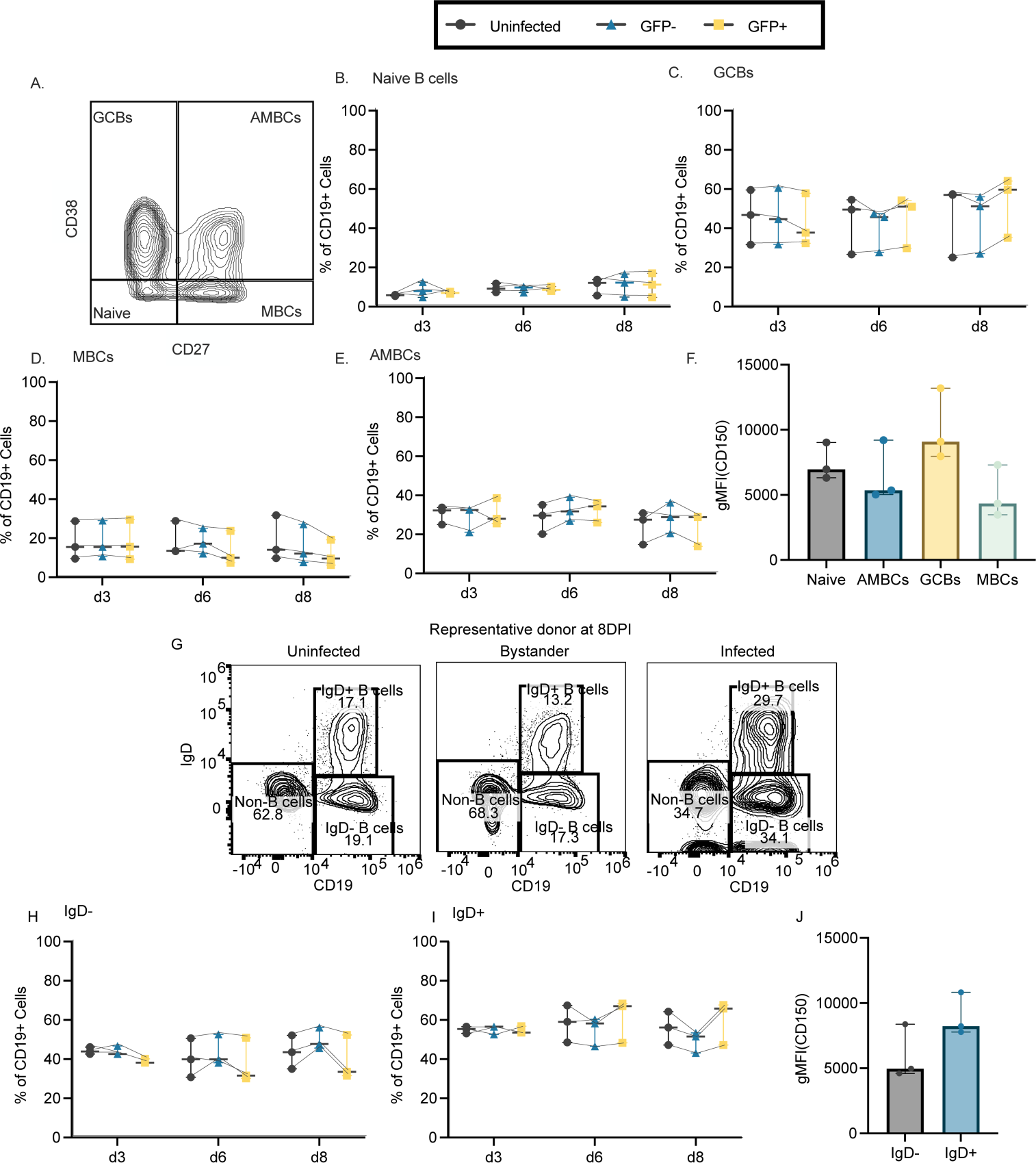
MeV infects B cell subsets proportionally. Donors (n=3) from Figure 3 were immunophenotyped to identify B cell subsets using CD38 and CD27 within the CD19^+^ cells, with the gating strategy utilized shown in **(A)**. Naive (CD27^-^CD38^-^; **B**), germinal center (CD27^-^CD38^-^; **C**), memory (CD27^+^CD38^-^; **D**), and activated memory (CD27^+^CD38^+^; **E**) B cells were quantified by comparing their frequency among both uninfected and GFP^+^ cells over time. CD150 expression was calculated for each population among uninfected cells at 6 days post-infection and the mean fluorescent intensity (MFI) of CD150 expression is shown **(F)**. Representative flow plots are shown demonstrating the enrichment of IgD^+^ B cells among all infected cells at 8DPI **(G)**. Susceptibility to infection was assessed by quantifying the frequency of IgD-**(H)** or IgD^+^ **(I)** cells among uninfected, bystander, or infected cells. CD150 expression on IGD- and IgD^+^ populations at day 6 are shown in **(J)**. For all immunophenotyping panels, significance was determined by two-way ANOVA using the Geisser-Greenhouse correction with Tukey’s multiple comparison test. For panel **(F)** significance was determined by one-way ANOVA using Friedman’s test with Dunnett’s multiple comparison test. Significance in **(J)** was determined using the Wilcoxon matched-pairs signed rank test. For all plots, the median with the 95% confidence interval is shown.

**Figure S5 (related to Figure 3):**
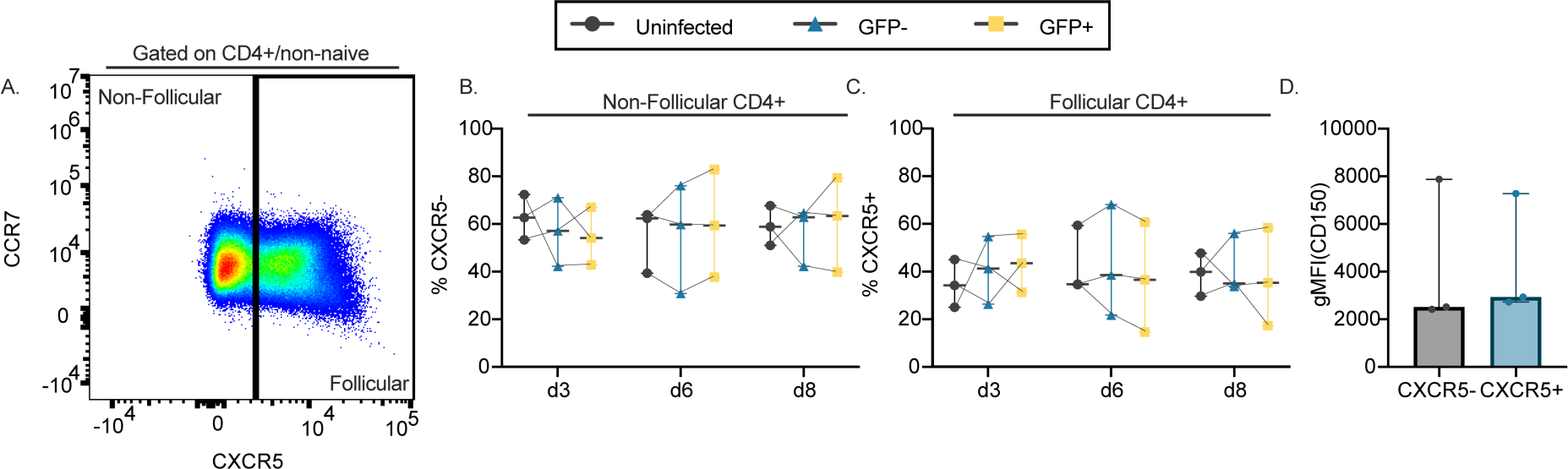
CXCR5 status has no impact on the susceptibility of CD4^+^ cells. Non-Naïve CD4^+^ cells (n=3) were subset based on CXCR5 status as shown in (**A**). Shown are the frequency of non-follicular (CXCR5^-^; **B**) and follicular (CXCR5^+^; **C**) cells among uninfected, GFP^-^ (bystander), or GFP^+^ non-naïve CD4^+^ cells. CD150 expression was compared between non-follicular and follicular cells (**D**). Significance for **B-C** was determined by two-way ANOVA using the Geisser-Greenhouse correction with Tukey’s multiple comparison test, and CD150 expression significance was determined by the Wilcoxon matched-pairs signed rank test (**D)**. For all plots, the median with the 95% confidence interval are shown.

**Figure S6 (related to Figures 2 and 4):**
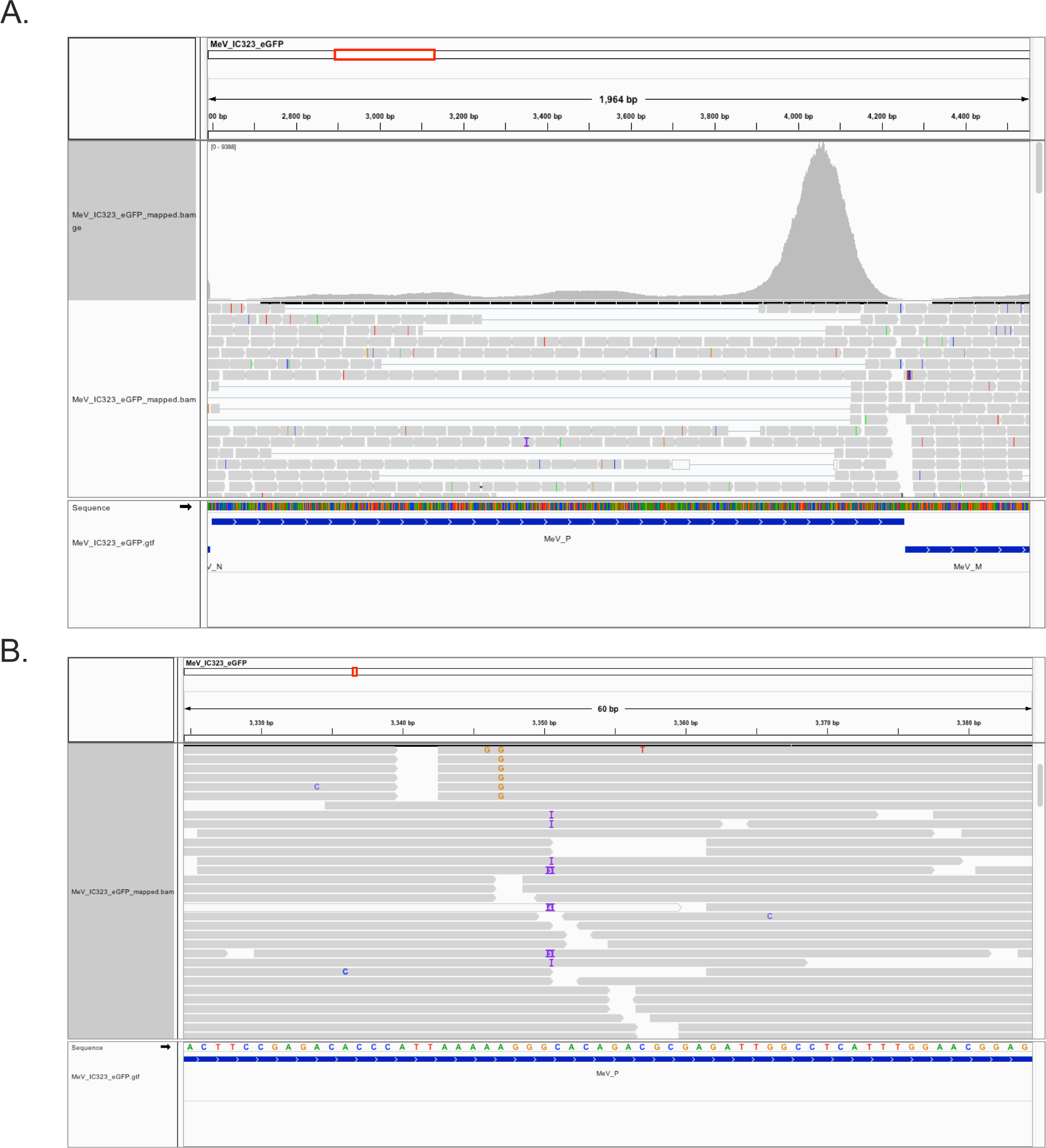
Detection of P-editing by scRNA-Seq. **(A)** Read coverage for all libraries along the length of the MeV P transcript. The red box denotes the region of the MeV linear genome that is shown in the histogram box, and the boxes under the histogram are a selection of reads that map to these positions. Gray bars indicate that the read matches the reference sequence, with colored letters representing mismatches to the reference. **(B)** Zoomed-in coverage of the p-edited region, showing representative reads that map to this region of the gene in gray. “I” represents indel mappings, which may indicate P-edited transcripts. There were 235 total reads covering the putative edit site.

**Figure S7 (Related to Figure 5).**
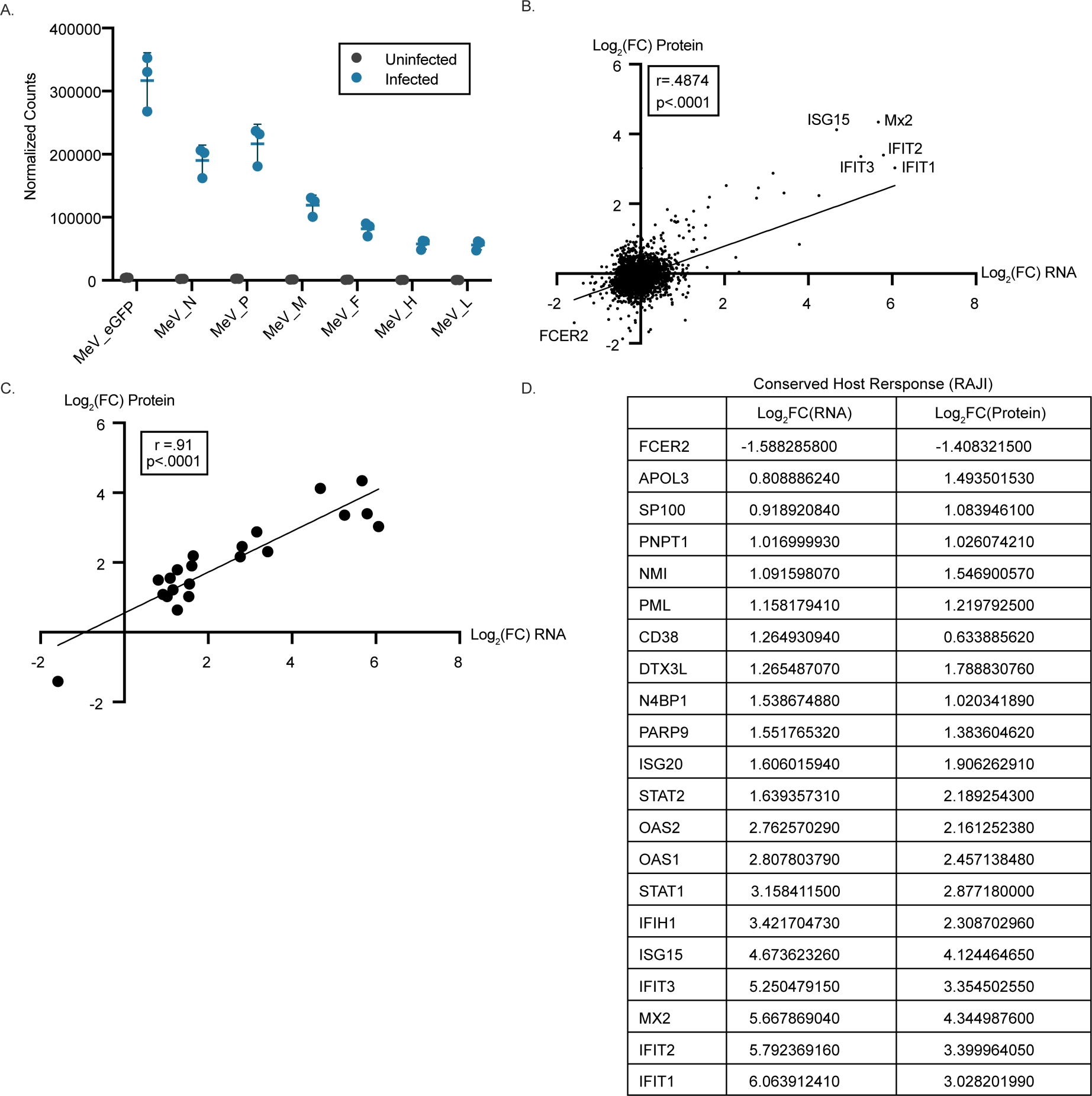
Comparison of the transcriptome and proteome in infected Raji cells. Raji cells were infected with MeV and collected for bulk RNA sequencing (n=3). **(A)** Detection of MeV transcripts in infected cells is shown as normalized counts for each MeV gene compared to uninfected controls. Median and 95% confidence intervals are shown. **(B)** Correlation plot showing the relationship between differentially expressed proteins (y-axis) identified by MS in Figure 5 with differentially expressed transcripts identified by bulk RNA sequencing (x-axis). Log2FC values for transcripts and proteins that were detected in both RNA and protein assays are shown. Simple linear regression was conducted, and the Pearson correlation value was reported on the plot, along with the significance of the correlation. **(C)** Correlation plot demonstrating the relationship between the list of significantly altered proteins (y-axis) and transcripts (x-axis). Hits that were significant in both assays are denoted in the table shown in **(D)**.

